# Comparative venom analysis between melanistic and normally-colored phenotypes of the common adder (*Vipera berus*)

**DOI:** 10.1101/2024.03.18.585504

**Authors:** Lennart Schulte, Lilien Uhrig, Johanna Eichberg, Michael Schwartze, Ingve Auth, Miriam Schulz, Thomas Lindner, Paul Hien, Kornelia Hardes, Andreas Vilcinskas, Tim Lüddecke

**Affiliations:** Institute for Insect Biotechnology, Justus Liebig University Giessen, Heinrich-Buff-Ring 58, 35392 Giessen, Germany; Department of Bioresources, Fraunhofer Institute for Molecular Biology and Applied Ecology, Ohlebergsweg 12, 35392 Giessen, Germany; LOEWE-Centre for Translational Biodiversity Genomics, Senckenberganlage 25, 60325 Frankfurt, Germany; Institute for Landscape Ecology, University of Münster, GEO 1, Heisenbergstraße 2, 48149 Münster, Germany; Institute for Zoology and Evolutionary Biology, University of Regensburg, Universitätsstraße 31, 93053 Regensburg, Germany; Natural History Museum – Leibniz Institute for Evolution and Biodiversity Science, Invalidenstraße 43, 10115 Berlin, Germany; Venture for Interconnection, Protection, Education and Research in Adders e.V. (VIPERA), Brunnenplatz 9, 92355 Velburg, Germany

## Abstract

Snake venom is an ecologically-relevant functional trait directly linked with a snake’s fitness and survival, facilitating predation and defense. Snake venom variation occurs at all taxonomic levels, but the study of variation at intraspecific level is still in its early stages. The common adder (*Vipera berus*) exhibits considerable variation in color phenotypes across its distribution range. Melanistic (fully black) individuals are the subject of myths and fairytales, and in German folklore such “hell adders” are considered more toxic than their normally-colored conspecifics. However, these claims have never been formally tested. Here, we provide the first comparative analysis of venoms from melanistic and normally-colored common adders. Specifically, we compared the venom profiles by SDS-PAGE and RP-HPLC, and tested the protease, phospholipase A_2_ and cytotoxic activities of the venoms of both phenotypes. Phospholipase A_2_ activity was similar in both phenotypes, whereas general protease activity was higher in the melanistic venom, which was also more cytotoxic at two concentrations (6.25 and 12.5 µg/ml). These minor differences between the venoms of melanistic and normally-colored adders are unlikely to be of clinical relevance in the context of human envenomation. In light of our results, the claim that melanistic adders produce more toxic venom than their normally-colored conspecifics appears rooted entirely in folklore.

## 2 Introduction

Venom has evolved multiple times independently in the animal kingdom, and serves the three major functions of hunting, defense, and intraspecific competition (Casewell et al., 2013; Fry et al., 2009; Schendel et al., 2019). Concerning snakes, evidence suggests that snake venom evolved to mainly aid predation (Barlow et al., 2009; Daltry et al., 1996; Holding et al., 2021). However, selection on snake venom composition resulting from it being used for defensive purposes has also been demonstrated (Kazandjian et al., 2021). Snake venoms are commonly defined as complex mixtures of biomolecules, described as toxins, most of which are proteins or peptides (Casewell et al., 2013). Initially, snake venoms were considered to be well-defined and conserved within species and between closely related species. However, several studies have revealed the dynamic nature of this ecologically-relevant trait, highlighting both interspecific (Damm et al., 2021) and intraspecific differences in venom composition (Casewell et al., 2020). In the latter case, venom profiles have been shown to differ according to sex (Menezes et al., 2006), life stage (Cipriani et al., 2017) and geographic origin (Zancolli et al., 2019). Although this phenomenon of venom variation provides insight into the evolutionary ecology of snakes and other venomous animals, it also makes the treatment of envenomation more challenging (Casewell et al., 2020). If the symptoms of envenomation differ according to the individual animal, this affects both clinical decision making and the efficacy of any anti-venom that is administered (Casewell et al., 2020). Venom variation is particularly important in snakes because snakebite is classified by the World Health Organization as a priority 1 neglected tropical disease (Chippaux, 2017). Up to 2.7 million cases are estimated to occur every year, leading to hundreds of thousands of fatalities and/or permanent disabilities (Gutiérrez et al., 2017; Kasturiratne et al., 2008).

In Europe, the most widespread venomous snake is the common adder (*Vipera berus*). The taxonomy of this species is not completely resolved, with several subspecies being the subject of ongoing discussion (Geniez, 2018; Oskyrko et al., 2024; Speybroeck et al., 2016, 2020). Adult specimens are on average 50–70 cm in length, rarely exceeding 80 cm (Mallow et al., 2003). The neonates feed primarily on small amphibians and reptiles, although adults will also prey on birds and mammals (Mallow et al., 2003; Otte et al., 2020; Samsonov et al., 2022). One characteristic trait of common adders is their wide range of color and pattern phenotypes (Mallow et al., 2003; Otte et al., 2020). The most common phenotype is a pale grey (males) or brown (females) base color with a dark dorsal zig-zag pattern that forms a V-shaped motif on the head. These normally colored individuals are often also referred to as “cryptic”, however some authors argue for an aposematic function of this normal coloration (Forsman, 1995; Martínez-Freiría et al., 2017, 2020; Valkonen et al., 2011; Wüster et al., 2004). The dorsal zigzag pattern varies in width, and the body color can shift towards reddish brown or even blue. The most unique phenotypes are copper-colored and black (melanistic) specimens. Based on our own observations, their phenotype can change over the course of maturation. For instance, most of the melanistic adders are born with a cryptic pattern and change to the melanistic phenotype, while relatives of the same litter retain a cryptic phenotype. Interestingly, color phenotypes may have ecological repercussions in adders. For example, melanistic adders occur in many populations but vary in abundance even though they are easier for predators to see (Andrén and Nilson, 1981; Martínez-Freiría et al., 2020). The fitness trade-off may involve the higher sunlight-to-body-temperature conversion rate of melanistic compared to cryptic forms, particularly in colder temperatures and during shorter days, helping to increase metabolic activity and foraging, which could be translated into earlier maturation, an earlier start to the annual mating season, shorter egg development times, and a better fitness overall (Andrén and Nilson, 1981; Forsman, 1995; Martínez-Freiría et al., 2020). A selection of color phenotypes of the common adder is presented in Figure 1.

**Figure 1.**
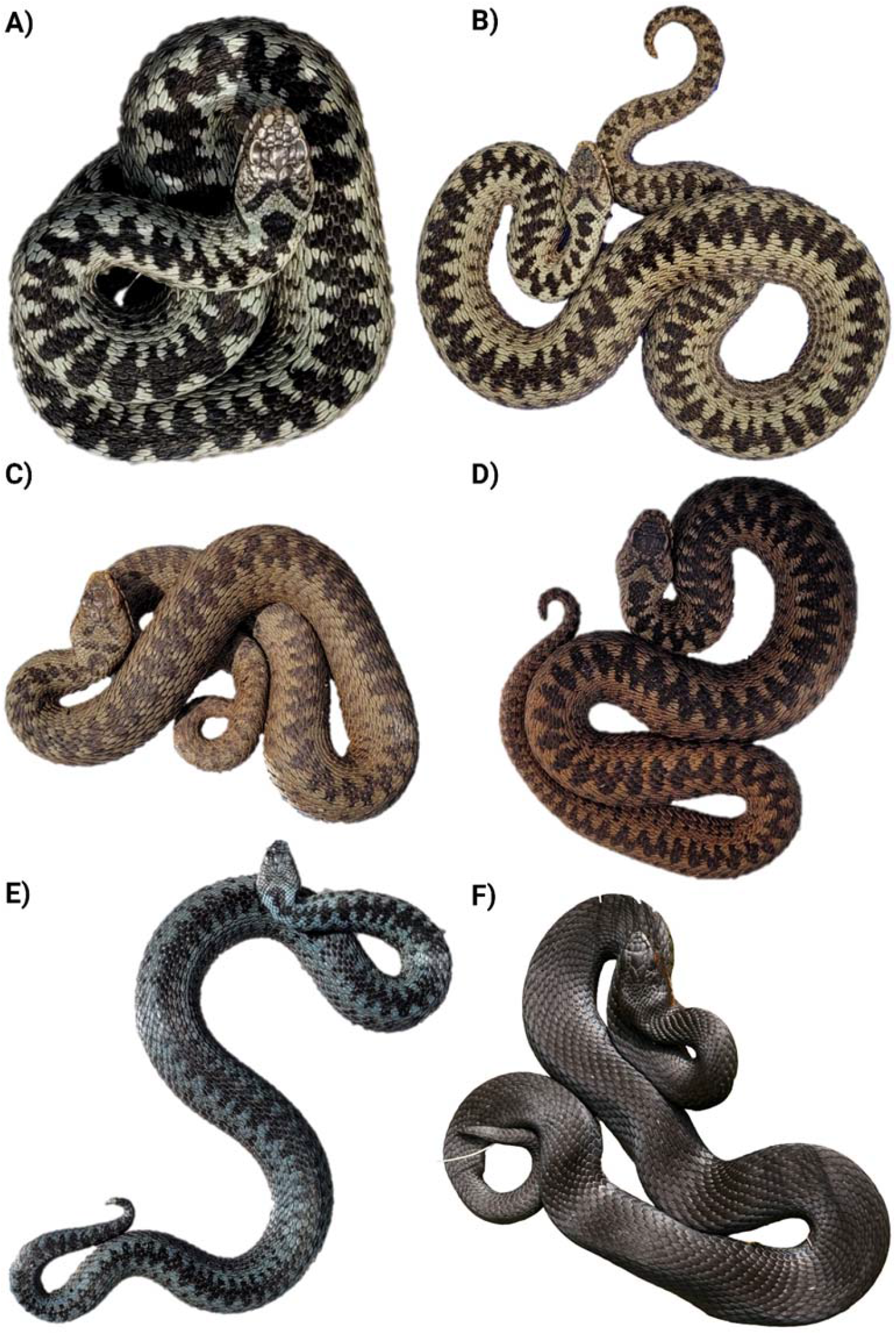
Color and dorsal pattern variation in German *Vipera berus*: A) specimen with classic color phenotype, exhibiting grayish background coloration, conspicuous black dorsal zig-zag pattern, and a clear V-shaped head marking; B-D) specimens exhibiting red/brown/copper background coloration and different levels of intensity in the dorsal pattern and V-shaped marking; E) specimen with blueish background coloration, less marked zig-zag pattern, and inconspicuous V-shaped head marking; F) melanistic specimen, with fully black coloration and unnoticeable dorsal pattern and V-shaped head marking.

Compared to other European vipers, human envenomation by *V. berus* is rarely fatal, but the incidence is high because the species is broadly distributed, including colder regions that are not inhabited by other venomous snakes (Di Nicola et al., 2021; Hermansen et al., 2019; Paolino et al., 2020). Due to this high bite incidence, the common adder ranks among the clinically most relevant venomous snakes snakes (Di Nicola et al., 2021; Hermansen et al., 2019; Paolino et al., 2020). The venom profile of *V. berus* has been analyzed (in a few cases using modern venomics technologies) and seems to match the typical viperine profile dominated by members of the phospholipase A_2_ (PLA_2_) family, snake venom metalloproteases (svMPs), serine proteinases (svSPs) and C-type lectins (CTLs), including snaclecs and C-type lectin-related proteins (Al-Shekhadat et al., 2019; Bocian et al., 2016; Latinović et al., 2016). *V. berus* also appears to show intraspecific venom variation because the proportion and abundance of dominant and lesser toxin families varies between studies and according to the origin (Damm et al., 2021) and assigned subspecies (Damm et al., 2024) of each specimen.

Interestingly, the common adder and particularly its venom are mentioned in the European folklore, especially across German speaking countries. Here, melanistic adders have a reputation for greater aggression and toxicity than adders exhibiting lighter color phenotypes, and were initially considered a separate species, earning the name “hell adder” (Blum, 1888; Otte et al., 2020). This largely reflects superstitions connected to dark animals, including cats, dogs and ravens, which have been integrated into myths and folklore, often as symbols of bad luck, treachery, evil and witchcraft (Byghan, 2020). While it is now understood that ‘hell adders’ are simply melanistic animals, in certain parts of Europe, such as the more rural parts of Germany, black-colored adders are still feared as more toxic in the general public, despite lacking scientific basis. Phenotype-dependent venom variation is rarely investigated (Avella et al., 2023), but a comparison of venom from a rare melanistic rattlesnake (*Crotalus durissus terrificus*) and its conspecifics provided some support for the hypothesis (da Silva et al., 1999). However, a single specimen does not provide enough data to draw firm conclusions.

Here we set out to clarify whether or not melanistic adders may indeed have distinct venom profiles or increased toxicity in comparison to normally colored conspecifics. Therefore, we carried out the first comparative analysis of venom composition and venom bioactivity in adders of different phenotypes to. We compared the venom profiles, enzymatic activities and cytotoxicity of venom from normally colored (cryptic, CRY) and melanistic (MEL) phenotypes. Our work provides insights into venom variation in adders and could be used as the basis for future comparative venomics studies in snakes and to facilitate adder conservation.

## 3 Materials and Methods

### 3.1 Venom

Venom was donated by members of the German Society of Herpetology and Herpetoculture, and was sourced from captive individuals of German origin. Venom was pooled from nine adult males of each phenotype (MEL or CRY) and lyophilized. Only male individuals of German origin were used for this study, to rule out potential sex or locality-based effects. Lyophilized venoms were redissolved in double-distilled water and aliquots were stored at – 20 °C.

### 3.2 Compositional venom profiling

Compositional venom profiling was carried out as previously described (Schulte et al., 2023) by combining reducing and non-reducing sodium dodecyl-sulfate polyacrylamide gel electrophoresis (SDS-PAGE) with 5 µg of each sample and reversed-phase high-performance liquid chromatography (RP-HPLC) with 125 µg of each sample. Chromatograms (280 nm) were normalized against an initial equilibration run to reduce noise. Raw gel image is provided in Supplementary Figure S1.

### 3.3 Phospholipase A_2_ assay

Phospholipase activity was measured using the EnzChek Phospholipase A_2_ Assay Kit (Invitrogen, cat. no. E10217) for 96-well plates. Venoms were redissolved in 1x Reaction Buffer and added in triplicates at final concentrations of 3.125, 6.25, 12.5, 25 and 50 µg/ml. Autolytic activity was minimized by handling all samples on ice and reducing the pre-incubation time. After incubation for 30 min at room temperature, the plates were transferred to a Synergy H4 Hybrid Microplate Reader (BioTek) and analyzed using Gen 5 v2.09 software (BioTek). The signal was detected at 515 nm following excitation at 470 nm. Values were averaged and normalized against the positive control (5 U/ml purified bee venom phospholipase A_2_, 100%) and negative control (1x Reaction Buffer, 0%). Normalized raw data from the phospholipase A_2_ activity assays are provided in Supplementary Table S1.

### 3.4 Protease activity assay

We used a non-specific Protease Activity Assay Kit (Calbiochem, cat. no. 539125) for 96-well plates as previously described (Schulte et al., 2023). Venoms were redissolved in double-distilled water and added in triplicates at final concentrations of 25, 50, 100, 200 and 400 µg/ml, as described above. The reactions were incubated for 2 h at 37 °C, shaking at 120 rpm on a Multitron device (Infors HT) and the OD_492_ was measured in an Eon microplate reader (BioTek). The signals were averaged and normalized to the positive control (166 µg/ml trypsin, 100%) and negative control (double-distilled water, 0%). Normalized raw data from the protease activity assay are provided in Supplementary Table S2.

### 3.5 Cell viability assay

The cytotoxicity of venoms was assessed in Madin-Darby canine kidney II (MDCK II) cells using the CellTiter-Glo Luminescent Cell Viability Assay (Promega, cat. no. G7570) as previously reported (Hurka et al., 2022). The venoms were redissolved in cultivation medium and added to final concentrations of 1.56, 3.125, 6.25, 12.5 and 25 µg/ml. Luminescence was measured using a Synergy H4 Hybrid Microplate Reader, and mean values for each treatment were normalized against the positive control (100 µM ionomycin, 0% growth) and negative control (cultivation medium, 100% growth). Normalized raw data from the cell viability assay are provided in Supplementary Table S3.

### 3.6 Hemolytic assay

The hemolytic activity was based on a previously described method (Sæbø et al., 2023) that was adjusted to a 96-well format. To purify horse erythrocytes, whole blood was washed in Alsever buffer until the supernatant prepared by centrifugation (804 *g*, 5 min, 4 °C) was clear. The pellet was resuspended using a cut 1000-µl tip to reduce shear forces. Purified erythrocytes were diluted in Alsever buffer to a final concentration of 1% (w/v). Venom was redissolved to final assay concentrations of 5, 10, 20, 40 and 80 µg/ml and added in triplicate to the erythrocyte suspension at a 1:1 ratio in V-bottom 96-well plates. The signal was detected at OD_405_ and mean values were normalized to the positive control (1% Triton X-100, 100% lysis) and negative control (Alsever buffer, 0% lysis). Normalized raw data from the hemolytic activity assay are provided in Supplementary Table S4.

## 4 Results

### 4.1 Compositional profiling

We fractionated 5 µg of each venom by reducing SDS-PAGE (Figure SDS, A), resulting in protein denaturation and the separation of monomeric proteins, and also by non-reducing SDS-PAGE (Figure SDS, B), to preserve any disulfide bonds holding together multimeric complexes. This provided a comprehensive overview of the protein composition to facilitate our analysis of putative toxin families. Reducing SDS-PAGE revealed protein mass ranges of 12–70 kDa for both phenotypes. MEL venom featured slightly more intense bands at ∼14, ∼23, ∼31 and 47–70 kDa, with a distinct band at ∼55 kDa (Figure SDS, A). Non-reducing SDS-PAGE revealed a mass range of 12–115 kDa for both phenotypes. MEL venom featured more intense bands at 76–80, ∼90 and ∼115 kDa compared to CRY venom, but the latter featured a more intense band at 32–34 kDa (Figure SDS, B).

**Figure 2.**
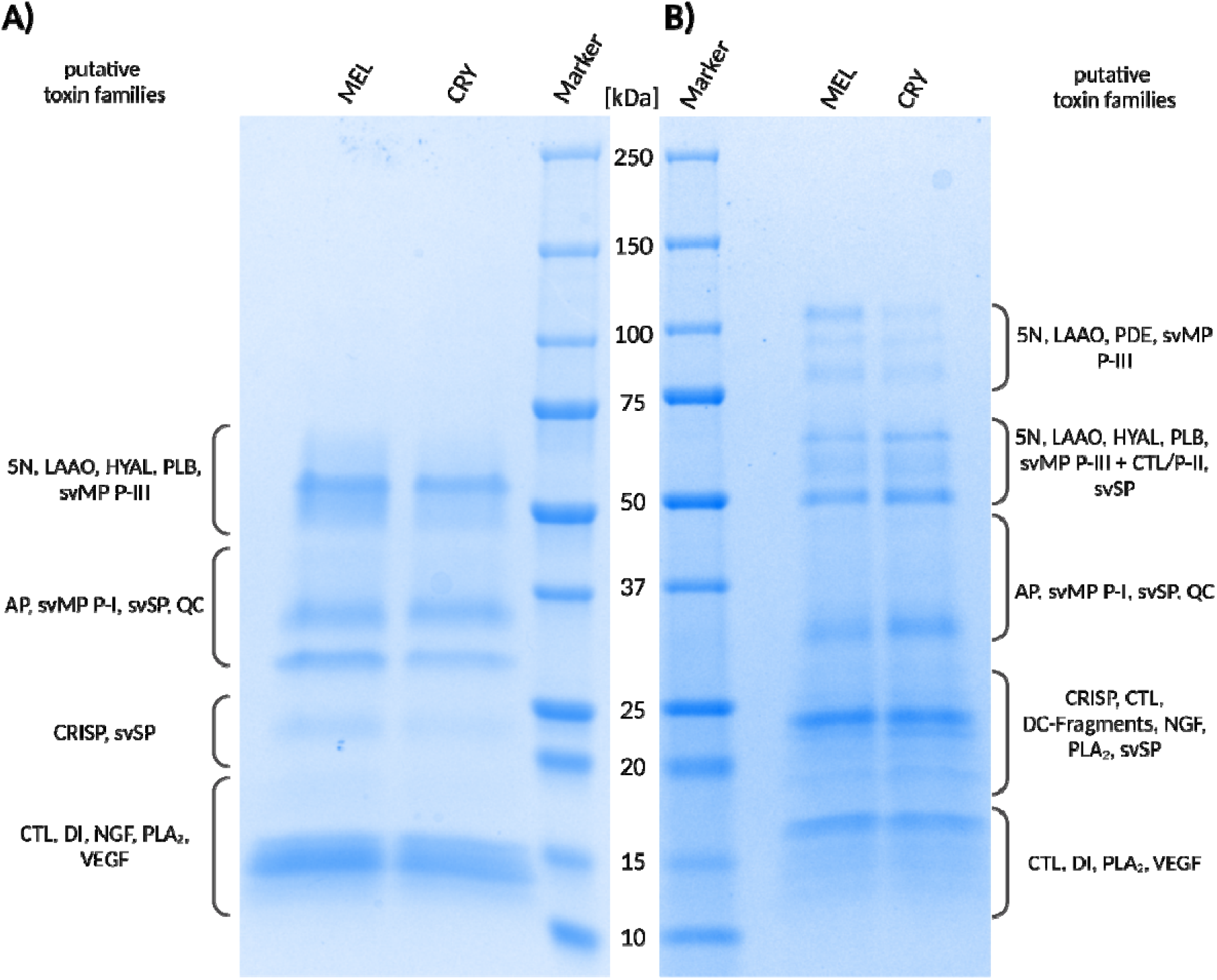
Analysis of *Vipera berus* venom samples from melanistic (MEL) and cryptic (CRY) phenotypes by SDS-PAGE under (A) reducing and (B) non-reducing conditions. Putative toxin families were predicted according to described Viperidae toxin families, their polymerization potential, and the previously reported composition of *V. berus* venom (Al-Shekhadat et al., 2019; Bocian et al., 2016; Damm et al., 2024; Latinović et al., 2016). Abbreviations: 5N, 5′-nucleotidase; AP, aminopeptidase; CTL, C-type lectin including snaclec and C-type lectin-related proteins; DC-Fragments, Disintegrin-like/cysteine-rich protein (fragments); HYAL, hyaluronidase; LAAO, l-amino acid oxidase; NGF, nerve-growth factor; PLA_2_, phospholipase A_2_; PLB, phospholipase B-like; PDE, phosphodiesterase; QC, glutaminyl cyclase; svMP, snake venom metalloprotease; svSP, snake venom serine proteinase; VEGF, vascular endothelial growth factor.

We then separated 125 µg of each venom by RP-HPLC (Figure HPLC) revealing peaks over a retention time of 2–27 min, with the highest peak in both samples at ∼9 min. Clusters of peaks were observed for both venoms at 17:30–18, 19:30–21, 21:30–23 and 24:30–27 min, although the peaks in the last three clusters were smaller in MEL than CRY venom. The smaller peaks and peak cluster were mostly similar in appearance, but the small peak cluster at 15:30 min was much less apparent in CRY venom.

**Figure 3.**
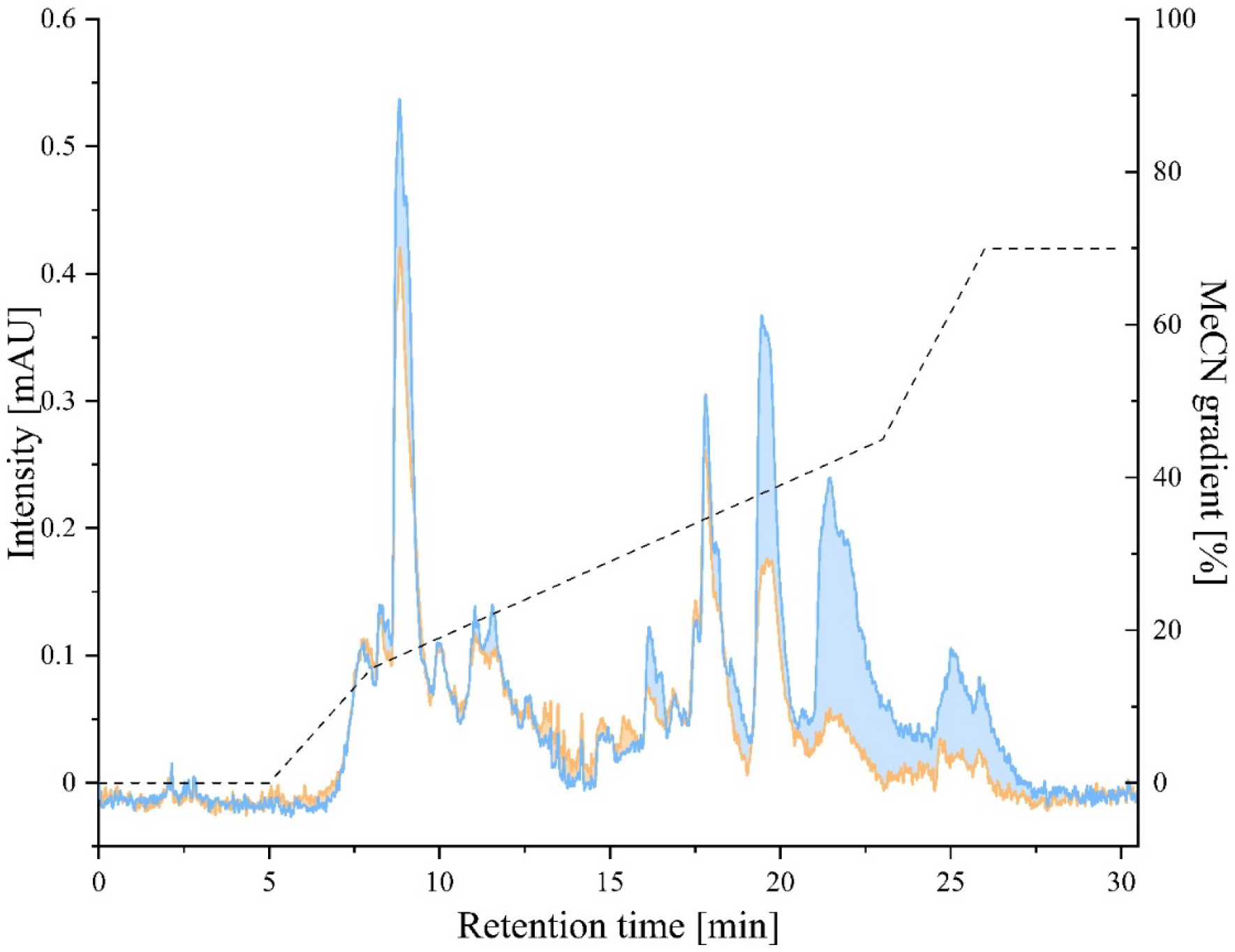
Chromatograms of *Vipera berus* venoms. The chromatograms (280 nm) correspond to venoms obtained from melanistic (orange line) and cryptic (blue line) specimens, eluted and measured using a gradient of acetonitrile (dashed line) over a 30 min time frame. Differences in peak intensities are highlighted by coloring the area between the traces, using the color code of the venom exhibiting the more intense signals.

### 4.2 Bioactivity profiling

Next, we tested the bioactivity of the venoms in enzymatic assays covering the dominant venom toxin families of *V. berus* (PLA_2_ and the svMP and svSP families) and in cytotoxicity assays against a canine somatic cell line and equine erythrocytes.

PLA_2_ activity (Figure PLA_2_) in both venoms was assessed at concentrations of 3.125, 6.25, 12.5, 25 and 50 µg/ml. The observed activity ranged between 38.86% (CRY, 3.125 µg/ml) and 101.47% (MEL, 50 µg/ml), relative to assay buffer (0%) and 5 U/ml bee venom PLA_2_ (100%). PLA_2_ activity was similar between MEL venom (3.125 µg/ml, 41.87%; 6.25 µg/ml, 62.33%; 12.5 µg/ml, 81.06%; 25 µg/ml, 101.37%; 50 µg/ml, 101.47%) and CRY venom (3.125 µg/ml, 38.86%; 6.25 µg/ml, 64.32%; 12.5 µg/ml, 85.15%; 25 µg/ml, 98.09%; 50 µg/ml, 99.76%), with standard deviations exceeding relative activity disparities at most concentrations.

**Figure 4.**
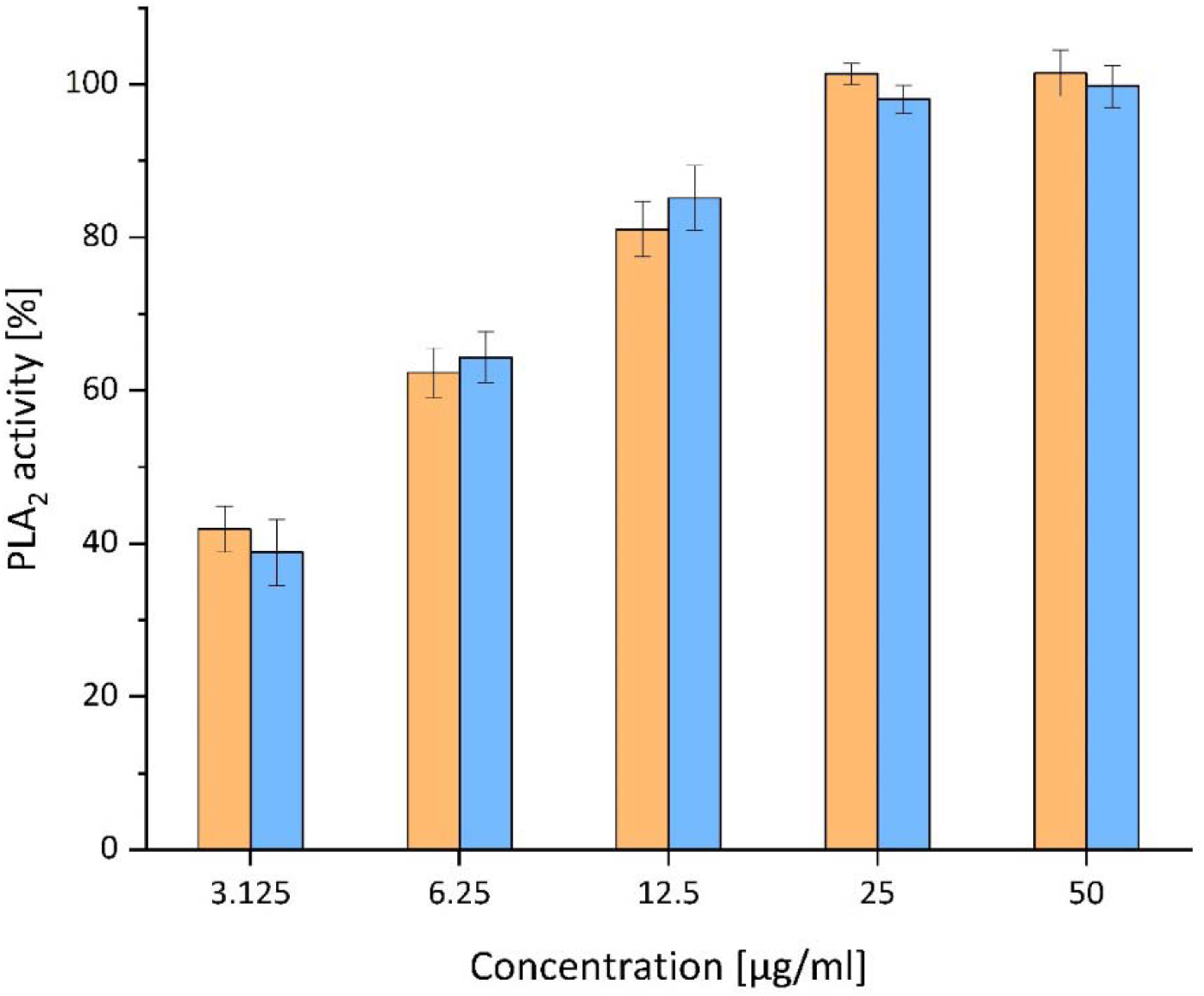
Phospholipase A_2_ (PLA_2_) activity of the two *Vipera berus* venoms. Venoms from melanistic and cryptic specimens are represented in orange and blue, respectively. The graph represents the results of the five concentrations tested (i.e., 3.125, 6.25, 12.5, 25, and 50 µg/ml). Values were normalized against purified bee venom PLA_2_ (positive control, 100%) and buffer (negative control, 0%). Data are means ± standard deviations (n = 3).

In the protease activity assay, we applied the MEL and CRY venoms at concentrations of 25, 50, 100, 200 and 400 µg/ml. The protease activity (Figure PAA) ranged between 1.57% (CRY, 25 µg/ml) and 46.00% (MEL, 400 µg/ml) relative to the negative control (0%) and the positive control (100%). At every tested concentration, the MEL venom (25 µg/ml, 7.46%; 50 µg/ml, 10.80%; 100 µg/ml, 22.49%; 200 µg/ml, 33.83%; 400 µg/ml, 46.00%) showed higher activity than the CRY venom (25 µg/ml, 1.57%; 50 µg/ml, 3.41%; 100 µg/ml, 8.21%; 200 µg/ml, 15.60%; 400 µg/ml, 28.18%). Thus, the MEL venom showed a clear tendency for higher protease activity compared to CRY venom.

**Figure 5.**
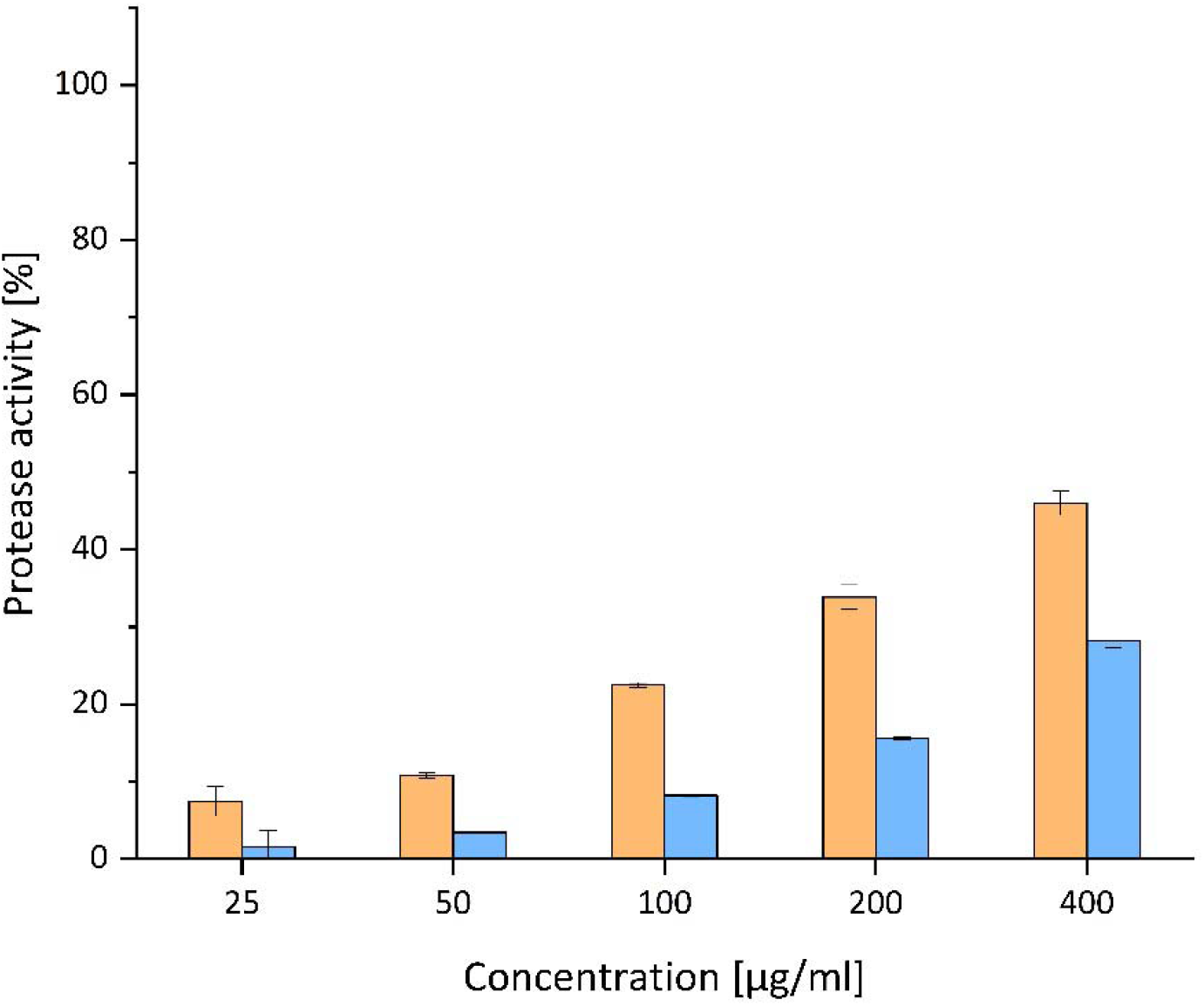
Protease activity of the two *Vipera berus* venoms. Venoms from melanistic and cryptic specimens are represented in orange and blue, respectively. The graph represents the results of the five concentrations tested (i.e., 25, 50, 100, 200, and 400 µg/ml). Values were normalized against purified trypsin (positive control, 100%) and ddH_2_O (negative control, 0%). Data are means ± standard deviations (n = 3).

The cytotoxic effects of the venoms were tested at concentrations of 1.56, 3.125, 6.25, 12.5 and 25 µg/ml against canine MDCKII cells (Figure Cytotox). Strong cytotoxicity was observed regardless of the venom type at a concentration of 25 µg/ml (MEL 99.51%, CRY 99.65%), but no effects were detected at 3.125 µg/ml (MEL –1.50%, CRY –24.55%) or 1.56 µg/ml (MEL –15.96%, CRY –19.04%). At 6.25 µg/ml (MEL 13.73%, CRY –14.26%) and 12.5 µg/ml (MEL 72.46%, CRY 32.24%), the cytotoxicity of the venoms from melanistic animals was higher than that of the CRY venom, although again the samples size was insufficient for statistical analysis.

**Figure 6.**
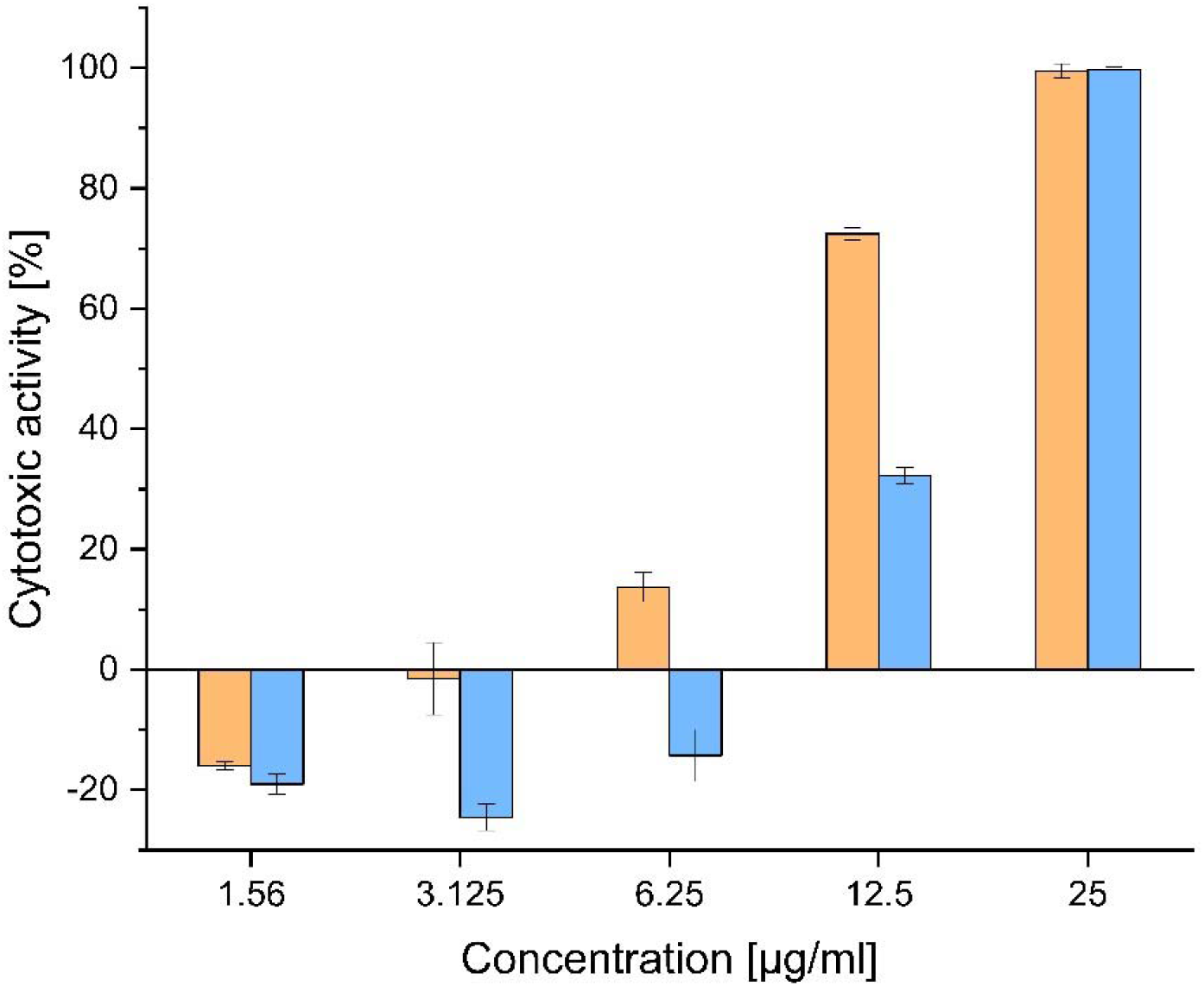
Normalized cytotoxicity of the two *Vipera berus* venoms. Venoms from melanistic and cryptic specimens are represented in orange and blue, respectively. The graph represents the results of the five concentrations tested (i.e., 1.56, 3.125, 6.25, 12.5, and 25 µg/ml) against MDCKII cell lines. Data are means ± standard deviations (n = 3).

We also tested the venom against purified horse erythrocytes at concentrations of 5, 10, 20, 40 and 80 µg/ml (Supplementary Figure S), but no hemolysis was detected regardless of the venom type or its concentration (Supplementary Figure S4).

## 5 Discussion

Melanistic common adders have a reputation across Europe for being more toxic than normally-colored ones. Although this perception appears to be based on folklore and superstition rather than empirical evidence, it was never tested scientifically. To our knowledge, this is the first work formally investigating the presence of differences between the venoms of specimens of the two phenotypes in terms of composition and biological activities.

The comparison of CRY and MEL by SDS-PAGE and RP-HPLC venom profiles revealed qualitative similarities (i.e., number of bands and peaks did not differ between the two venoms) but quantitative differences (i.e., bands and peaks presented different intensities between the two venoms). This variation partly translated into differences in enzymatic activity among the dominant toxin families, with MEL venom showing a trend for higher protease (svMP and svSP) activity, whereas PLA_2_ activity was comparable between the samples. However, we observed little difference in cytotoxicity between the venoms when tested against a canine (MDCKII) cell line, and there was no hemolytic activity against equine erythrocytes.

The analysis of the venom profiles through SDS-PAGE at reducing and non-reducing conditions allowed us to identify putative venom toxins based on known Viperidae venom components, their potential to form multimeric complexes, and the previously reported venom composition of *V. berus* (Al-Shekhadat et al., 2019; Bocian et al., 2016; Damm et al., 2024; Latinović et al., 2016). The MEL banding pattern was generally more intense than its CRY counterpart, especially at ∼31 and 47–70 kDa in reducing gels and at 76–115 kDa in non-reducing gels. Based on the analysis by REF, these bands likely represent dominant *V. berus* venom components such as svMP and svSP, suggesting that they are more abundant in the MEL venom. Previous assessments of *V. berus* venom proteases showed strong activity for fractions in the range 50–66 kDa in reducing gels (Malina et al., 2017). In the same size range of this proteolytically active fraction, the bands were more intense in MEL venom, potentially explaining differences observed in the bioactivity profiling.

Indeed, although the analysis of enzyme activity reflecting the dominant components of *V. berus* venom revealed no consistent differences in PLA_2_ activity, it suggested an increased protease activity in MEL venoms. Significant differences in protease activity between individual *V. berus* specimens but consistent levels of PLA_2_ activity have been reported before (Malina et al., 2017). In light of this, it is possible that the composition and activity of *V. berus* venom could vary in terms of proteases, but remain relatively stable in the context of PLA_2_ activity within populations and between individuals and phenotypes.

The cytotoxicity of the MEL venom was higher than that of the CRY venom, but only when tested at 6.25 or 12.5 µg/ml. This limited effect in a narrow concentration range is unlikely to be clinically relevant because adult adders inject 10–18 mg of venom during human envenomation (Al-Shekhadat et al., 2019). State-of-the-art *in vivo* assays to assess the potency of venom usually test for necrosis, lethality, neurotoxicity and other effects in mice or rats which is difficult to translate into human symptomatics (Calderón et al., 1993; Maievskyi and Slieptsova, 2023; Malina et al., 2017). Similarly, our cytotoxicity data cannot directly predict the symptoms of envenomation or the lethal dosage. That said, it is a useful approximation towards the potential for degenerative necrosis in the affected area, with differences occurring in a limited range of concentrations. Further, we observed no evidence of hemolytic activity against equine erythrocytes at the concentrations we tested. Case reports mention hemoconcentration, which is often caused by hemolytic activities, as symptoms of *V. berus* envenomation in only 5% and 10% of incidents, respectively, suggesting hemolysis as a minor effect (Hermansen et al., 2019).

The analysis of intraspecific venom variation is relatively novel in the field of toxinology, and the role of a snake’s phenotype in its occurrence has rarely been considered relevant thus far, resulting in a lack of data. However, a rare case of melanism in the pit viper *C. d. terrificus* prompted a comparison of its venom profile with a normally-colored specimen (da Silva et al., 1999). The analysis of protein bands revealed a core venom composition, albeit with some quantitative variation, as well as unique bands in each specimen. Although the analysis of a single specimen is not representative, the study nevertheless showed a similar trend to our observations, with the venom from the melanistic snake yielding more intense bands corresponding to larger toxins. However, the venom composition of *V. berus berus* specimens from Russia (Al-Shekhadat et al., 2019; Latinović et al., 2016) and the Slovakian Republic (Bocian et al., 2016), as well as one *V. berus barani* specimen from Türkiye (Damm et al., 2024), revealed remarkable quantitative diversity for PLA_2_ (10% (Latinović et al., 2016), 17.9% (Damm et al., 2024), 25.3 % (Al-Shekhadat et al., 2019), 59% (Bocian et al., 2016)), as well as svMP (0.2% (Damm et al., 2024), 3.15% (Bocian et al., 2016), 17.2% (Al-Shekhadat et al., 2019), 19% (Latinović et al., 2016)) and svSP (15% (Bocian et al., 2016), 16.2% (Al-Shekhadat et al., 2019), 31% (Latinović et al., 2016), 46.1% (Damm et al., 2024)). In relation to this extreme compositional variation across the sampling sites in previous studies, the venom variation we observed by SDS-PAGE and RP-HPLC was marginal and only affected the abundance of certain fractions rather than their overall diversity. Also, recent studies in related species of the subfamily Viperinae suggest ontogenetic shifts, diet and environmental conditions as important drivers of venom variation, which are certainly more dominant than phenotype dependencies (Adamude et al., 2023; Avella et al., 2023, 2022; Casewell et al., 2009; op den Brouw et al., 2021). Previous studies of intra-population venom variation in *V. berus* from Hungary revealed individual venom variation similar in magnitude to our observations, with quantitative as well as qualitative differences (Malina et al., 2017). Accordingly, the few quantitative differences suggested by our SDS-PAGE and RP-HPLC profiling as well as the functional differences determined in our bioassays, may represent only the normal biological variability of *V. berus* venom instead of an attribute of a specific color phenotype.

Considering the apparently great extent of biological variation within *V. berus* and given the many factors that can influence venom composition, future studies should be based on larger sample sizes. In addition, they should account for relevant confounding factors such as the life stage, sex and diet of the specimens, the population, season, and probably even individual variations. This will allow the statistical validation of bioactivity assays to definitively address the conundrum of phenotype-dependent venom variation.

## 6 Conclusion

Several color phenotypes of common adders (*Vipera berus*) are known in Europe. Particularly, the melanistic specimens are subject to myth and folklore, where they are said to be of higher toxicity than normally colored conspecifics. Here we provide the first compositional and functional determination of phenotype-dependent venom variation in common adders, using melanistic and normally-colored individuals as a model system. By rigorous implementation of chemical profiling methods (SDS-PAGE and RP-HPLC) we unveiled, that venoms of both phenotypes contain fundamentally the same components yet not necessary at the same quantities. As for functional differences, which we investigated via *in vitro* bioassays targeting important viperine venom activities, we detected some differences. Venoms from melanistic specimens seem to display higher protease activity and higher cytotoxicity, albeit only at a narrow concentration range. On a first glance, these results support a conceptual difference between venoms of both phenotypes. However, as they only correspond to a few factors tested and, especially for the cytotoxicity assays, are only detected at lower concentrations they are unlikely to be of clinical relevance. Considering the tremendous extent of venom variation reported in *V. berus* across its distribution range, the differences observed in our experiments may represent the normal biological variability within this species instead of a trait of melanistic animals. We recommend further investigations of that topic using larger sample sizes and additionally assays to fully resolve this question. However, interpreting our data in light this known venom variability, the limited range of experiments returning significant differences and the magnitude of differences measured, it seems that the reputation of MEL phenotypes is based on human experience of envenomation and is probably an irrational superstition after all.

## 7 Supplementary material

The Supplementary Material for this article can be found in Figure S1 and Table S1-S4

## 9 Acknowledgements

We thank the members of the German Society for Herpetology and Herpetoculture for donating the venom samples used in this study. Further, we are grateful to Ignazio Avella, Maik Damm and Till Röthig for comments made on an earlier version of this manuscript. A.V. acknowledges generous funding from the Hesse Ministry of Science and Art (HMWK) via the LOEWE Centre for Translational Biodiversity genomics. T.L. received funding from the German Research Foundation (DFG) through the project FunVen (Project ID 505696476). MDCKII cell line was kindly provided by Eva Böttcher-Friebertshäuser, Institute of Virology, Philipps University, Marburg. We are grateful to Richard M. Twyman for language editing. Figures 1 and 2 were created using BioRender under the following licenses: RF26KPWQKD (Figure 1), BV26JJZF3Q (Figure 2)

