## Supplementary Figure S1 for "Comparative venom analysis between melanistic and normally-colored phenotypes of the common adder (*Vipera berus*)"

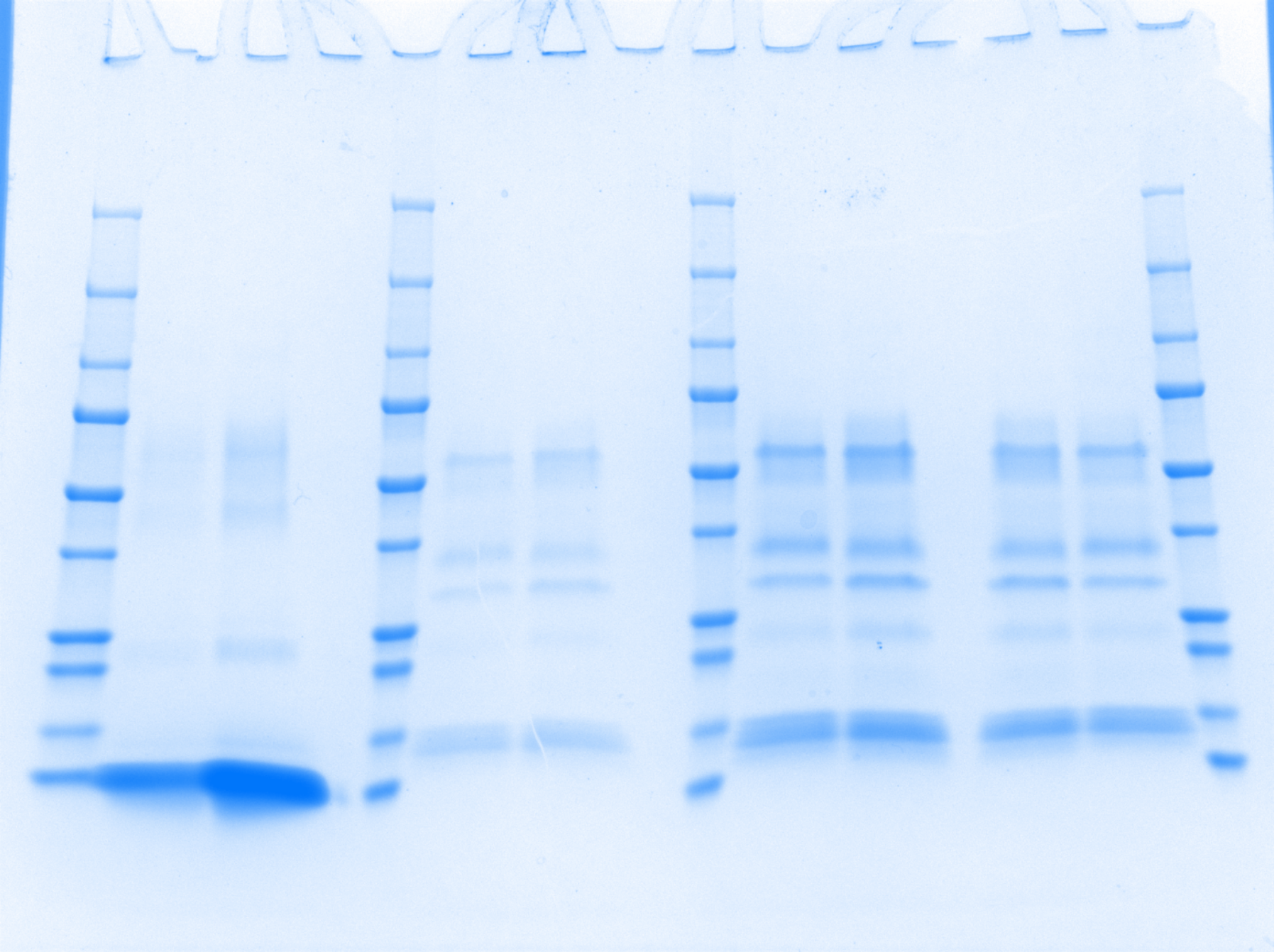

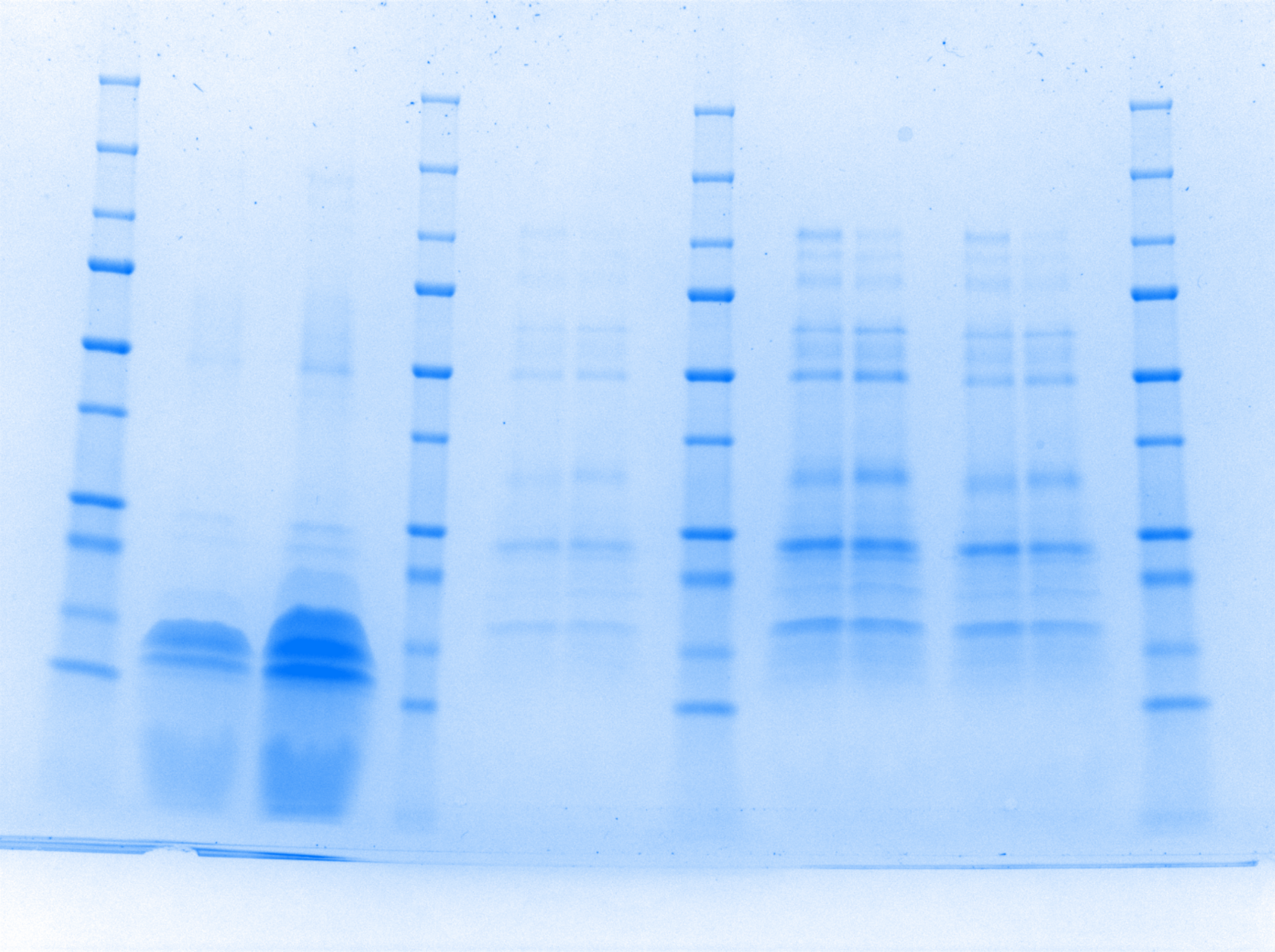

MEL

CRY

MEL

CRY

Marker

Marker

[kDa]

15

20

25

37

50

75

100

150

250

10

15

20

25

37

50

75

100

150

10

250

Supplemantary Figure S1: SDS-PAGE raw images of pooled Vipera berus venom samples from melanistic (MEL) and cryptic (CRY) phenotypes under A) reducing and B) non-reducing conditions. Non-relevant lines, left and right of the section of interest, are cropped out.
